# Identifying behavioral links to neural dynamics of multifiber photometry recordings in a mouse social behavior network

**DOI:** 10.1101/2023.12.25.573308

**Authors:** Yibo Chen, Jonathan Chien, Bing Dai, Dayu Lin, Zhe Sage Chen

**Author notes:** (D.L.) and (Z.S.C.). Equal contributions (Y.C. and J.C.).

## Abstract

Distributed hypothalamic-midbrain neural circuits orchestrate complex behavioral responses during social interactions. How population-averaged neural activity measured by multi-fiber photometry (MFP) for calcium fluorescence signals correlates with social behaviors is a fundamental question. We propose a state-space analysis framework to characterize mouse MFP data based on dynamic latent variable models, which include continuous-state linear dynamical system (LDS) and discrete-state hidden semi-Markov model (HSMM). We validate these models on extensive MFP recordings during aggressive and mating behaviors in male-male and male-female interactions, respectively. Our results show that these models are capable of capturing both temporal behavioral structure and associated neural states. Overall, these analysis approaches provide an unbiased strategy to examine neural dynamics underlying social behaviors and reveals mechanistic insights into the relevant networks.

## INTRODUCTION

Social interactions involving aggression, defense and mating are known to involve a large number of distributed brain areas, among which the hypothalamic-midbrain neural circuits play an especially important role (Anderson, 2016; Falkner et al., 2016; Lo et al., 2019; Lischinsky and Lin, 2020; Lovett-Barron et al., 2020; Liu et al., 2022; Wei et al., 2023; Guo et al., 2023; Mei et al., 2023a, 2023b). During aggressive behaviors, the brain uses multidimensional signals to coordinate discrete actions (Falkner et al., 2020). However, a more comprehensive understanding of the neural code underlying complex, naturalistic social behaviors remains an open goal in systems neuroscience. Recent technological advances in optical imaging have enabled the simultaneous and chronic recording of multiple brain regions in rodents, calling for the development of scalable and computationally efficient tools for large-scale data analysis (Chen and Pesaran, 2021). Multifiber photometry (MFP) is an in vivo calcium imaging method that detects average fluorescence intensity changes based on genetically encoded calcium (Ca^2+^) indicators and multimode optical fibers; MFP has been used to measure synchronous-firing neural population activity in multiple brain regions in freely-behaving animals (Cui et al. 2014; Kim et al., 2016; Meng et al., 2018; Sych et al., 2019). The lack of cellular resolution in MFP is counterbalanced by the ability to record from multiple subcortical regions of deep brain structures, or from molecularly-defined subpopulations with similar functions or homogeneous responses (Lin and Schnizer, 2016). Traditional strategies to dissect neural recordings often rely on prelabeled behavioral epochs defined by human observers or machine-based automated video analyses (Nilsson et al., 2020; Hsu and Yttri, 2021; Pereira et al. 2022), both of which may suffer from annotation inaccuracies due to camera occlusions or inter-rater inconsistency. Unsupervised learning may alleviate some of these challenges by instead identifying latent variables in neural data capable of meaningfully accounting for behavioral variabilities and common input (Liu et al., 2019; Calhoun et al., 2019; Pereira et al., 2020; Sani et al., 2021; Nair et al., 2023).

Mapping naturalistic and unconstrained behaviors to neural activity is a fundamental goal of neuroscience (Krakauer et al, 2017; Markowitz et al., 2018; Batty et al., 2019; Schneider et al., 2023). Latent variable models have been recently used to characterize complex, naturalistic aggressive behaviors (Aubry et al., 2022), or to characterize population calcium-imaging neuronal activity of the hypothalamus during aggressive behaviors (Nair et al., 2023). To the best of our knowledge, there has been no prior effort to apply these methods to MFP recordings during social behaviors, perhaps due to the ostensible challenge posed by limited temporal and cellular resolution in the face of complex naturalistic behaviors. Here we propose two latent state-space models, a continuous-state linear dynamical system (LDS) and a discrete-state hidden semi-Markov model (HSMM), to infer latent neural dynamics from MFP recordings of male mice during male-male and male-female social interaction. While the LDS approach has been widely used to characterize neural or joint neural-behavioral dynamics (Sani et al., 2021; Koh et al., 2023; Aghagolzadeh and Truccolo, 2016; Yu et al., 2009), the HSMM approach is a natural generalization of HMM analyses of neuronal spikes (Chen et al., 2014; Linderman et al., 2016; Liu et al., 2019), local field potentials (Cao et al., 2021), and calcium imaging (Tu et al., 2020). Since animal behavior is a phenotypic manifestation of well-orchestrated network activity, our goal is to uncover behaviorally relevant neural activity in the social behavior network (SBN). Our results suggest that MFP signals from the SBN can capture key temporal structures of aggressive and mating behaviors. In particular, the LDS approach identifies behavioral dynamics suggestive of behavioral subtypes, as well as behaviorally relevant latent dynamics that predict future neural observations and behavioral measures. The HSMM approach on the other hand can not only capture temporal structure and a hierarchy of aggressive behaviors but also discover micro-behavioral states that were sometimes missed by human observers. Together, these latent variable models uncover behavioral links to neural dynamics of population calcium signals in MFP recordings during mouse aggressive and mating behaviors.

## RESULTS

### Experimental overview

Using custom high-density multi-fiber arrays (**Figure 1A**), we recorded the Ca^2+^ activity of estrogen receptor type 1-expressing (*Esr1^+^*) populations from 13 brain regions in a well-defined SBN from freely-moving mice (Guo et al., 2023). Recordings were made in a male mouse as it socially interacted with a female or male mouse in a chamber (**Figure 1B** and **Methods**). The targeted SBN (**Figure 1C**) includes 5 hypothalamic regions: medial preoptic nucleus (MPN), anterior hypothalamic nucleus (AHN), ventrolateral part of the ventromedial hypothalamus (VMHvl), dorsomedial hypothalamus (DMH) and ventral premammillary nucleus (PMv); 5 amygdala regions: anterior medial amygdala (MeAa), posterodorsal medial amygdala (MeApd), posterior amygdala (PA), posteromedial cortical amygdala (CoApm) and posteromedial bed nucleus of stria terminalis (BNSTpm); and 3 other regions outside of amygdala and hypothalamus – ventral part of lateral septum (LSv), ventral subiculum (SUBv) and lateral periaqueductal gray (lPAG). The MFP recording appeared as simultaneously recorded 13 time series characterized by ΔF/F activity (**Figure 1D**) or deconvolved activity (**Figure 1E**).

**Figure 1.**
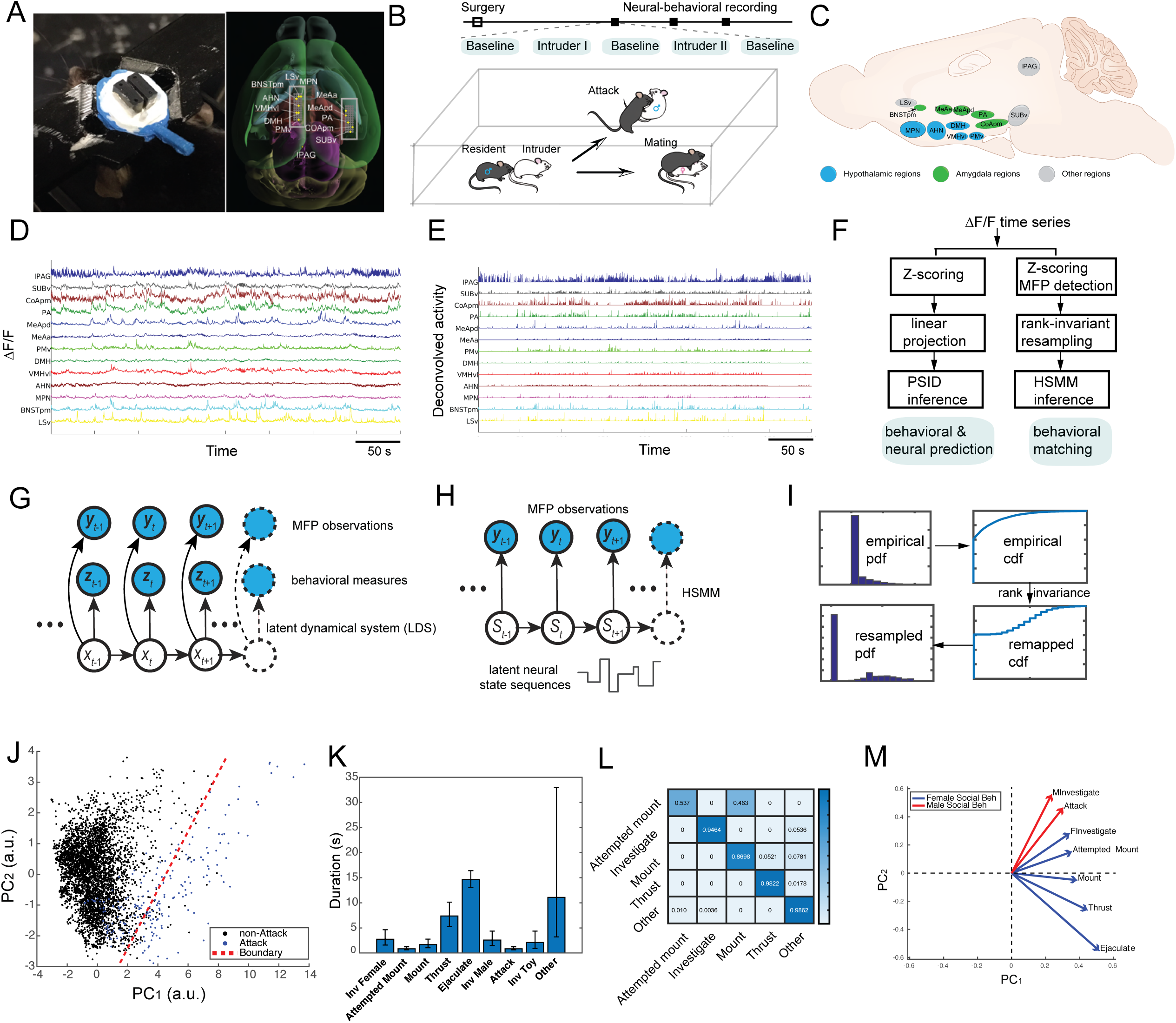
Experimental setup and joint neural and behavioral data analysis pipeline. (A) Schematic of MFP recording and optic fiber bundle. (B) Experimental protocol and recording timeline. (C) Illustration showing the recorded regions in mouse hypothalamus (blue), amygdala (green), and other brain areas (gray). (D) Representative snapshots of simultaneously recorded GCaMP6f trace (ΔF/F) of 13 brain regions in the mouse limbic system. (E) Deconvolved nonnegative activity from panel D. (F) Flowchart of neural data analysis pipeline. (G) Schematic of linear dynamical system (LDS) for inferring latent neural states that drive neural and behavioral measures. (H) Schematic of hidden semi-Markov model (HSMM) for inferring latent neural state sequences. (I) Illustration of rank-invariant resampling. (J) Principal component analysis (PCA) on selected 7-dimensional behavioral tracking measures (“*subj1_median_last_longest_hull”, “subj1_median_last_centroid_mvmt”, “subj1_median_last_tail_base_mvmt”, “both_mean_last_all_bp_mvmt”, “both_prctile_rank_mean_last_sum_bp_dist”, “subj2_shortest_last_median_bp_dist”, “subj1_prctile_rank_last_sum_centroid_mvmt”*) revealed a separation between “Attack” and “non-attack” behavioral episodes. Dotted line indicates the boundary inferred from generalized linear model (GLM) analysis. (K) Median duration of annotated behavior across all data. Error bar denotes 25% and 75% percentiles around the median. (L) Quantitative characterizations of annotated behaviors. (M) PCA showed two clusters of behaviorally relevant functional networks that explained the majority of variance of mean activity of 13 brain regions.

Upon collecting the MFP data from 13 regions from the mouse limbic system, we ran an analysis pipeline (**Figure 1F**) to compare neural dynamics with both continuous tracking-based behavioral features and discrete human-generated behavioral labels. Two latent variable models were used in our analysis pipeline. In the LDS approach (**Figure 1G**), we directly operated on the Z-scored ΔF/F activity and applied an established preferential subspace system identification (PSID) inference method to identify behaviorally relevant latent states (Sani et al., 2021). We focused our investigations on the relationship between MFP signals and 29 continuous behavioral features based on two-dimensional (2D) overhead tracking data of both subjects during male interaction behaviors. In the HSMM approach (**Figure 1H**), we operated on the inferred spiking proxy from Ca^2+^ activity (**Methods**) and further applied a rank-invariant resampling step to convert this activity into Poisson-distributed count data (**Figure 1I**); the results were further compared with human-annotated behavioral labels. In total, we analyzed 22 recording sessions collected from 5 male mice during the introduction of various male and female intruders (**Table S1**). However, only 16 recording sessions from 5 animals (#1, 6, 8, 12, 17) were accompanied by behavioral tracking data, which were analyzed using the LDS approach.

A total of 29 behavioral tracking features were generated from 2D tracking data; the features characterize physical variables associated with two interacting animals (pose, position, speed, distance, etc). During the annotated “*Attack*” epoch, principal component analysis (PCA) on selected tracking features (varying 7-29 dimensions) showed behaviorally meaningful separation (**Figure 1J**).

Additionally, human experimenters manually rated and classified the behavioral action states on a frame-by-frame basis. Animals’ behaviors were highly variable, and the number of annotated behavioral labels varied across sessions. Based on the behavioral labels, we quantified the mean event duration (**Figure 1K**) and empirical transition probabilities (**Figure 1L**)

Furthermore, we performed PCA on a “*region-by-annotated_label*” matrix, with each matrix element containing the mean ΔF/F activity averaged across time, sessions, and subjects within a specific annotated behavior. We found that the top two principal components (PCs) explained >95% variance, with male-male vs. male-female social behaviors separating in the loadings (**Figure 1M**).

### Joint neural-behavior analysis predicts latent dynamics of aggressive behavior

We first used an LDS to jointly model the Z-scored ΔF/F activity and behavioral tracking data. We assumed that the neural state and behavior were driven by a shared latent Gaussian-Markovian process, and the latent neural state consisted of both behaviorally relevant and irrelevant components (**Methods**). The model parameters were estimated based on a preferential subspace system identification (PSID) algorithm, and one-step-ahead neural/behavioral prediction was implemented by a recursive Kalman filter (Sani et al., 2021). We first used all MFP activity and selected seven behavior-tracking measures highly predictive of male aggressive attacks. We fit the model with the male-male interaction data (excluding male-female interaction epochs) for each session, followed by model testing on held-out data from the same session and other different sessions featuring the same animal.

In assessing the LDS-PSID approach on the cross-session data, we achieved a high degree of accuracy in neural observations, as seen in a very high correlation coefficient (CC) (**Figure 2A**). The hyperparameters such as the dimensionality of latent states were chosen by cross-validation. Generally, increasing the latent state dimension gradually increased the accuracy of neural prediction, but the accuracy of behavioral prediction saturated quickly (**Figure S1**). Notably, the model could generalize well not only within the same session, but also across sessions (**Figure 2B**). In contrast, the correlation between predicted and observed behavioral measures was lower, ranging from 0.2 to 0.7 depending on the specific behavior variable and specific session, and the model failed to capture faster dynamics or abrupt changes in behavioral tracking (**Figure 2C**). Additionally, the behavioral prediction accuracy was variable across individual behaviors and animals (**Figure 2D**). This issue may be due to (i) model limitations due to the linearity and stationarity assumptions; (ii) low fidelity or poor spatial resolution in the behavioral tracking data; or (iii) low temporal resolution of MFP signals and lack of specificity of the recorded brain regions accounting for behavioral tracking measures.

**Figure 2.**
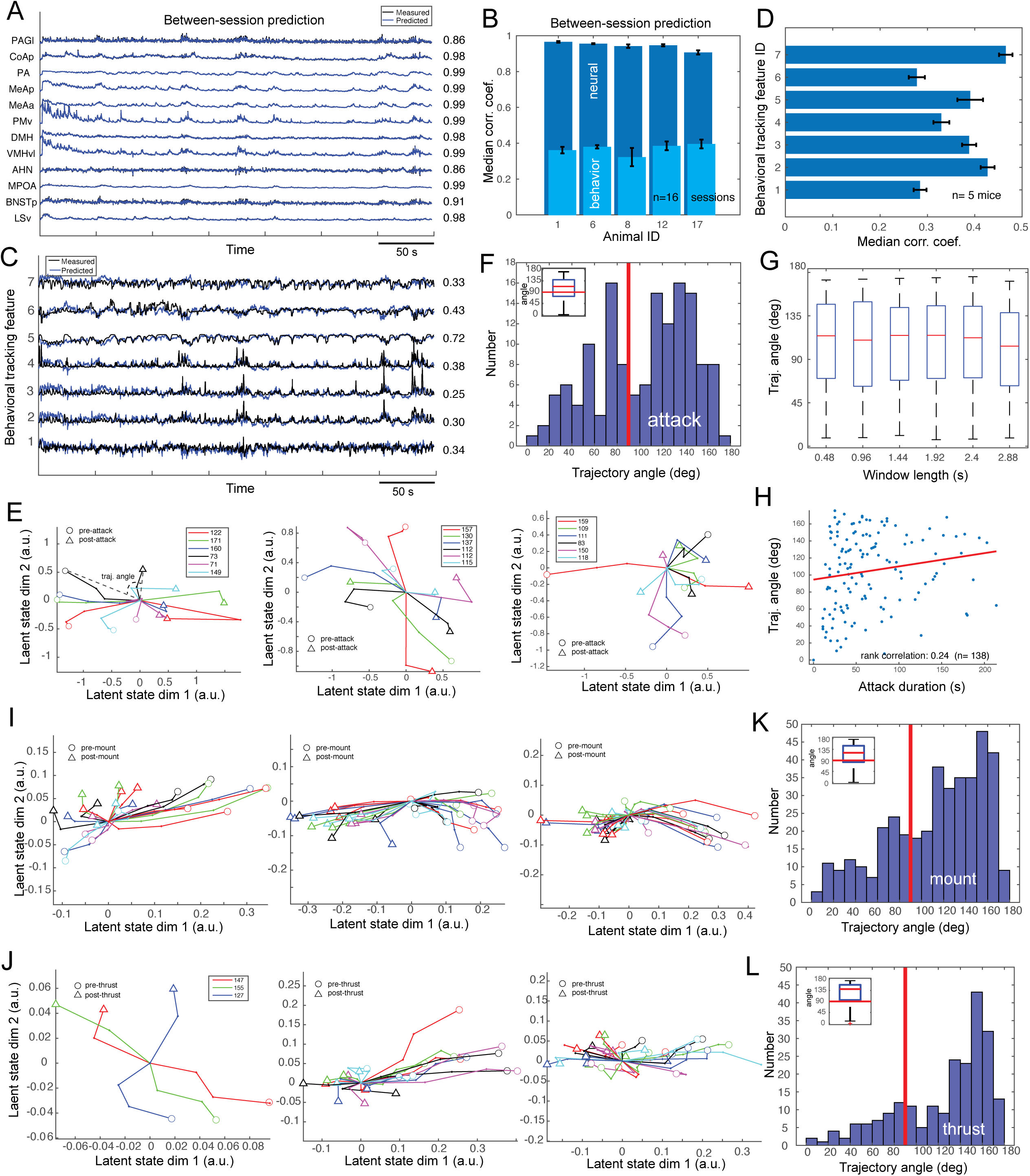
The LDS identify behaviorally relevant latent states. (A) Z-scored ΔF/F traces of 13 brain regions in MFP activity (black) and predicted traces (blue). Correlation coefficients between the measured and predicted traces were shown from the right. (B) Median correlation coefficients for between-session neural prediction (dark blue) and behavioral prediction (light blue). Error bars denote 25% and 75% percentiles across 16 sessions. (C) Selected 7 behavioral tracking traces (black) and prediction (blue) associated with panel A. Correlation coefficients between the measured and predicted traces were shown from the right. (Seven features #1-7 are “*subj1_median_last_longest_hull”, “subj1_median_last_centroid_mvmt”, “subj1_median_last_tail_base_mvmt”, “both_mean_last_all_bp_mvmt”, “both_prctile_rank_mean_last_sum_bp_dist”, “subj2_shortest_last_median_bp_dist”, “subj1_prctile_rank_last_sum_centroid_mvmt”)*. (D) Median correlation coefficients for between-session behavioral prediction across 7 behavioral measures. Error bar denotes IQR (25% and 75% percentiles) across 5 mice. (E) Visualization of aligned neural trajectories in “*Attack*” events from three representative recording sessions. 500-ms pre-attack and 500-ms post-attack trajectories are shown (start point: circles; end point: triangles). The angle from the start to end point define the trajectory angle. Figure legend denotes the angle in degree from the colored trajectories. (F) The distribution of trajectory angles among all attack events (n=138). Median angle was greater than 90 degrees (inset). (G) The median and distribution statistics of trajectory angle was robust with respect to the event window length (i.e., start-to-end-point duration). (H) The median trajectory angles had a positive correlation with respect to the attack event duration (Spearman’s rank correlation rho=0.24, *P* = 0.0039). Red line denotes a linear line fit. (I) Visualization of aligned neural trajectories in “*Mount*” events from three representative recording sessions. Figure legend same as panel E. (J) Visualization of aligned neural trajectories in “*Thrust*” events from three representative recording sessions (with one-to-one corresponding session in panel I). (K,L) The distribution of trajectory angles among all “Mount” (n=393) and “Thrust” (n=150) events. Median angle was greater than 90 degrees (inset).

To test the specificity of the 13 brain regions, we selected a subset of MFP activity from the predefined six regions of ABN and repeated the analysis; our results showed that the behavioral prediction accuracy further degraded (results not shown), suggesting that the unsatisfactory performance was partly due to insufficient spatial sampling in the mouse brain regions.

Furthermore, upon completion of inference we visualized the identified behaviorally relevant neural trajectories in low-dimensional latent space (**Methods**). Due to the lack of a clear trial structure, we aligned neural trajectories of a fixed duration, centered on the attack onset (**Figure 2E**). Interestingly, we found that the pre-attack trajectory tended to cross the origin and travel in a modified direction during the post-attack period, leading to a trajectory angle over 90 degrees among the majority of events (i.e., the angle between circles and triangles in **Figure 2E**), suggesting that many of these neural states were rotated in the state space (**Figure 2F**). Importantly, the median angle statistics were robust with respect to a wide range of duration parameters and consistently greater than 90 degrees (**Figure 2G**). Additionally, there was a significant correlation between the trajectory angles and the duration of attack events (Spearman’s rank correlation 𝜌=0.24, *P*=0.0039) (**Figure 2H**).

Next, we repeated similar analyses for “*Mount*” or “*Thrust*” events during male-female interactions. Surprisingly, we also observed very similar patterns for both “*Mount*” (**Figure 2I**) and “*Thrust*” (**Figure 2J**); the median angle statistics were around 135 degrees in both cases (**Figure 2K**). Together, these results suggested that the encoded neural patterns were rotated around the onset of aggressive and mating behaviors, yet there was a high degree of variability in neural trajectories across events.

### Cross-session prediction of aggressive behavior

Given good one-step-ahead prediction performance in the LDS-PSID approach, we further asked whether the onset of aggressive behavior (such as “*Attack*” during male-male interaction) can be reliably predicted based on MFP data. Because of the relatively small amount of attack sample points, we chose one animal with 5 recording sessions to demonstrate this predictive analysis. Specifically, we used the identified LDS model to first predict 13 neural or 20 joint neural-behavioral measures and then fed the predicted measures to the pretrained random forest (RF) or gradient boosting machine (GBM; Galar et al., 2001) classifier for detecting the target state on a rolling frame-by-frame basis (**Methods**). The leave-one-session-out analysis performances of these approaches were summarized in **Table 1**, where the mean AUC could reach 0.83-0.89, with a false detection alarm rate as low as 1.18 per min. Despite the limited amount of data, our results provide a proof-of-concept for online behavioral prediction.

**Table 1.** Summary of online detection performance for male attack behavior on a frame-by-frame basis (Animal #6). Leave-one-session-out results (mean ± SD, n=5) are shown (i.e., training on 4 sessions and testing on 1 session). N denotes the dimensionality of input features. The last column lists the ground truth attack rate for comparison with false alarm rate.

|  | AUC | Precision | F1-score | False alarm rate (per min) | Attack rate (per min) |
| --- | --- | --- | --- | --- | --- |
| Random forest (N=13) | 0.829±0.090 | 0.157±0.139 | 0.194±0.164 | 1.438±1.569 | 22.41±1.09 |
| Random forest (N=20) | 0.854±0.097 | 0.116±0.121 | 0.181±0.191 | 1.181±1.090 | |
| Gradient boosting machine (N=13) | 0.896±0.048 | 0.369±0.182 | 0.180±0.102 | 8.213±3.194 | |
| Gradient boosting machine (N=20) | 0.897±0.037 | 0.406±0.279 | 0.169±0.096 | 6.106±1.307 | |

### Unsupervised HSMM analysis uncovers behaviorally interpretable latent neural states

While the latent neural dynamics are continuous and potentially high dimensional, discrete characterization of neural dynamics may provide an alternative view in behavioral segmentation and labeling. We next employed a Bayesian nonparametric HSMM equipped with Monte Carlo sampling-based inference and automatic model selection to infer discrete latent neural states (Linderman et al., 2016). The HSMM generalizes the HMM by relaxing the assumption of history independence in the state transition, allowing more modeling flexibility (Chen et al., 2016).

Given the Z-scored MFP recordings (**Figure 3A**, top panel), we first detected the discrete event activity driving calcium signals from each region and further converted them to Poisson counting measures using a rank-invariant resampling method (**Figure 3A**, bottom panel), which enabled us to work with a conditional Poisson likelihood model accommodating efficient inference with conjugate priors (**Methods**). We fed the resampled observations of a complete MFP recording session to the HSMM and further inferred the latent state sequences in both training and held-out epochs, along with model parameters such as the firing rate matrix (**Figure 3B**, bottom panel).

**Figure 3.**
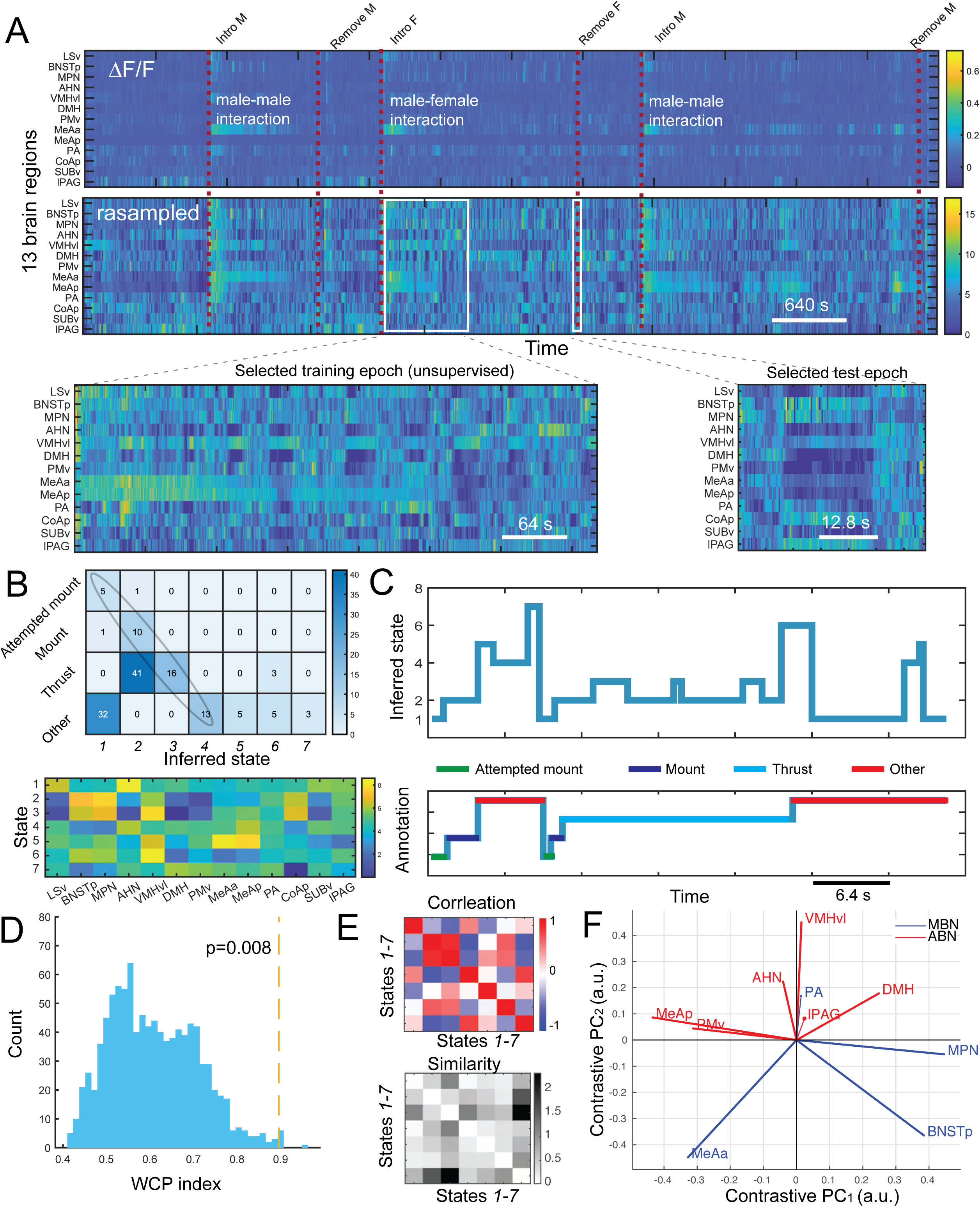
HSMM estimation results from a representative male-female interaction training epoch followed by testing on an unseen male-female interaction epoch. (A) Heatmaps of ΔF/F traces of 13 brain regions (top) in one recording session. Vertical red dashed line indicates the onset of annotated behaviors of Male (M) and Female (F) mice. Two white boxes represent the selected training and testing epochs of male-female interactions that share common annotated behaviors. (B) The derived confusion matrix (top) and state-firing rate matrix (bottom) from the selected test epoch example in panel A. The diagonal line of the confusion matrix indicates the degree of one-to-one correspondence between annotated behavior labels and inferred neural states. (C) Comparison of inferred state sequence (top) and annotated behavioral sequence (bottom) from the selected test epoch example in panel A. (D) Illustration of weighted column purity (WCP) index (red vertical dashed line) versus its shuffle distribution derived from the surrogate data. (E) Pairwise state correlation (top) and dissimilarity (bottom) matrices for the state-firing rate matrix in panel B. (F) By comparing the firing rate matrices during male-male and male-female interactions, contrast PCA roughly uncovered two functional networks: mating-biased network (MBN) and aggression-biased network (ABN).

Because of the behavioral diversity and naturalistic lack of imposed trial structure in each recording session, we conducted two types of analyses: (i) training and testing on behaviorally relevant epochs only, and (ii) training only but on the entire interaction (e.g., male-female or male-male interactions). Whereas this second type of analysis was straightforward and capable of using the whole dataset, the first type of analysis was only possible in some sessions when identical or similar behavioral labels were frequent enough to support segmentation into training and testing data.

In the first type of HSMM analysis, segmentation of MFP recordings into training and testing epochs was performed by a human analyst following several criteria as described below. For each session, we first identified the portions of the recording corresponding to an interaction with an intruder (there were 2-3 separate intruders per session, with intruders being a male, female, or toy). Each such portion (henceforth “*Interaction*”) ran from the first frame of the first annotation after the “Intro_M” and “Intro_F” annotations for male and female intruders, respectively, to the last frame of the last annotation preceding the “Rmv_M” and “Rmv_F” annotations, respectively (“Rmv” standing for “*Remove*”). We then excised from the interaction the *k* longest continuous periods annotated as “*Other*” to yield *j = k*+1 subsections, with the value of *k* swept across {1, 2, 3, 4, 5} for each interaction in each session for each subject. Each such set of *j* subsections produced for one value of *k* was termed a segmentation. Here we selected the value of *k* resulting in a segmentation that had at least two (out of *j*) subsections with good “duration” (i.e., number of timepoints in the subsection), “richness” (i.e., the proportion of timepoints in the subsection not labeled “*Other*”), and “diversity” (the proportion of all non-”*Other*” behaviors occurring in the entire interaction that occurred in the subsection). The subsection with the highest diversity score and longest duration (if there was a substantial difference among the subsections), in that order of importance, was designated as the training epoch. All remaining retained subsections were used as testing epochs, where the number of retained subsections varied between 2 and *j*. Note that the number of testing epochs was limited even if *j* > 2 because many subsections had low duration, richness, or diversity scores. By these criteria, we typically obtained 1 training epoch and 1-3 testing epochs for each recording session.

Upon completion of training, we matched behavioral labels and latent neural states (**Methods**) and then reordered the state identity to match the behavioral labels for better visualization. The inferred latent states had non-equal occupancy. In both training and post-hoc testing, we compared our neural state sequences with human-annotated behavioral labels and quantified the “column purity” from the confusion matrix (**Figure 3B**, top panel), which measures the specificity of a neural state for one behavioral label. In general, we found a consistent correspondence between neural states and annotated behaviors (**Figure 3C**) as suggested by a statistically significant “weighted column purity” (WCP) index (**Methods**) in the test epoch (Monte Carlo *P*<0.01, n=1000 random shuffles), where each annotated behavior was also weighted by its respective occupancy. In the male-female interaction example in **Figure 3A**, we obtained WCP statistics of 0.87 (**Figure 3D**). From the inferred state-firing rate matrix, we computed the pairwise Pearson’s correlation and dissimilarity based on Kullback-Leibler (KL) divergence (**Methods**). In the illustrated example in **Figure 3B**, the mean neural patterns of states 2 and 3 were similar (**Figure 3E**). In comparison to male-female interaction, behavioral transitions in male-male interaction were simpler. One representative example is shown in **Figure S2**.

From the *region-by-state* firing matrix, we further computed the region-by-region correlation matrix and applied contrastive principal component analysis (cPCA) (Abid et al., 2018) to find the subspace that distinguishes the patterns derived from two conditions (male-male vs. male-female interactions). We further found that the first two PC roughly separated two clusters of functional networks: a mating-biased network (MBN) and an aggression-biased network (ABN), where the MBN mainly consists of four regions: MPN, BNSTpm, PA, and MeAa, whereas the ABN mainly consists of six regions: VMHvl, PMv, MeApd, DMH, AHN and IPAG (Gao et al., 2023).

We next examined the source of row and column impurities in each test epoch. In the case of male-female interaction, some discrepancies between states and labels occurred during behavioral transitions (e.g., *“Attempted mount”*⤍*“Mount”* or “*Mount*”⤍“*Thrust*”), which could be due to discrepancies between human annotations and the HSMM estimation in identifying the boundary between two similar and naturally consecutive events. In addition, the HSMM tended to segment a longer duration into several short states (e.g., repeated transitions between “*Mount*” and “*Thrust*” in **Figure 3C**), whereas human annotations usually merged them into a single, long behavioral category (e.g., “*Thrust*” event in **Figure 3C**). Additionally, the same annotation label was often explained by more than one neural state (i.e., “row impurity”). However, this one-to-many mapping could be explained by the fact that these neural states tended to group under hierarchical clustering based on their state-firing rate similarity (**Figure 3B** and **Methods**). Further, low column impurity for each of these states indicates that they generally corresponded to only one behavioral annotation. Together, this suggests that these states may represent micro-behavioral categories or transition states. Our reasoning was also confirmed later by intensive computer simulations.

The second type of the HSMM analysis was similar, except that there was no separation between training and testing. This analysis was adopted to use a longer recording duration and test the model’s ability to deal with the predominant and highly heterogeneous “*Other*” category (during which the mice were not interacting) (**Figure 4A**). The performance was again characterized by the WCP index (for instance, the WCP was 0.918 in the illustrated example of **Figure 4A**). Since the “*Other*” behavioral label was heavily predominant in most recording sessions, many neural states could be assigned to capture the heterogeneity in this undefined “*Other*” category (**Figure 4B**). Based on the inferred state-firing rate matrix (**Figure 4C**), we clustered these states to reveal their relationship (**Figure 4D**). For instance, states 2 and 3 (matching approximately to Mount and Thrust, respectively) were very close. States 6, 9, 10 and 11 were also relatively similar in their activity patterns. These states predominantly occurred during undefined “*Other*” period with no social interaction, suggesting the possibility of micro-behaviors. Interestingly, the HSMM could sometimes identify mislabeling by human annotators. In the representative session in **Figure 4A**, we noticed the occurrence of states 1 and 2, which is often associated with “*Attempted mount” and “Mount*” sequence, although there was no such annotation in this case. When we re-inspected the video, we noticed a mounting event did occur but was originally omitted by the human annotator.

**Figure 4.**
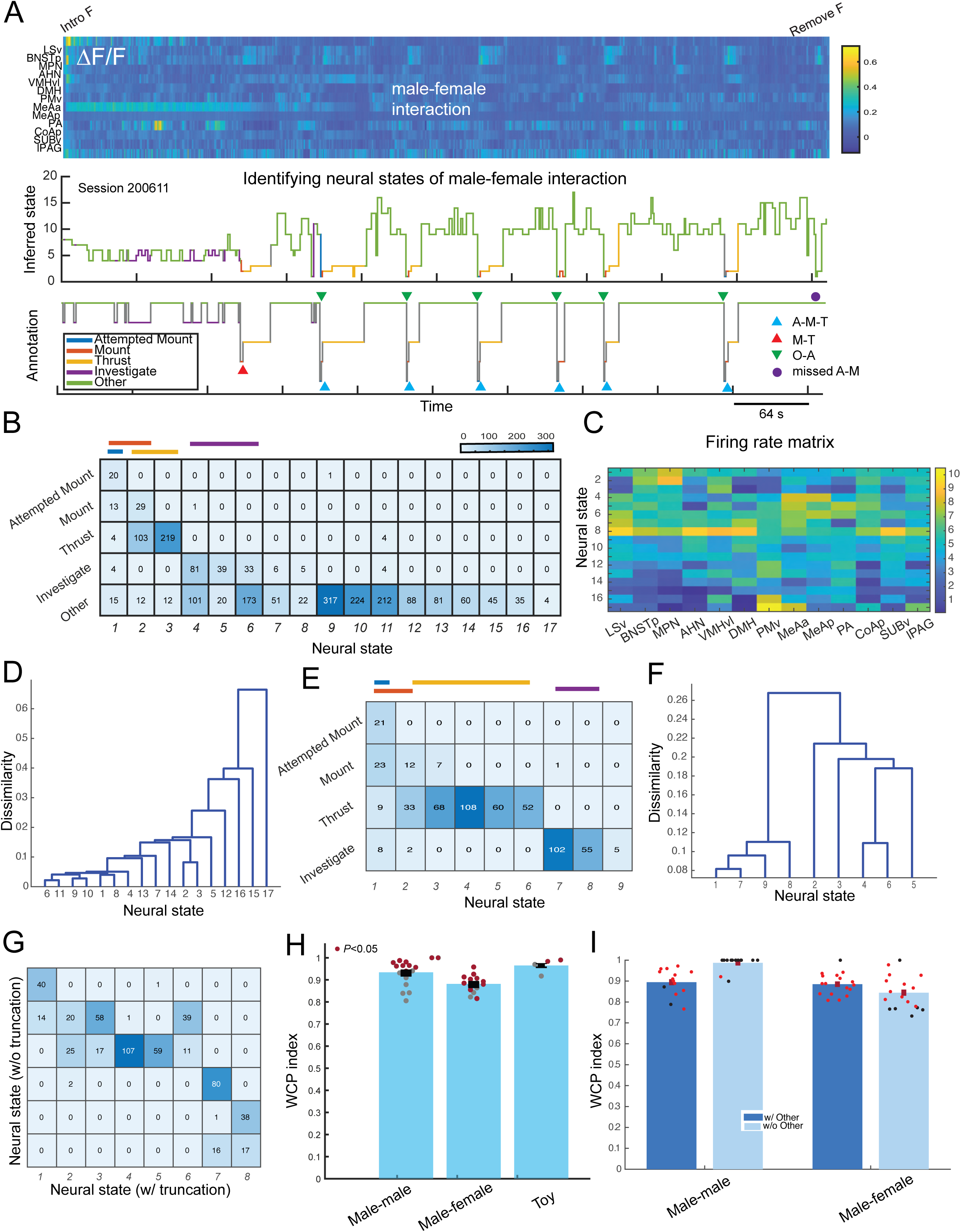
HSMM uncovers behaviorally interpretable latent neural states. (A) Top: Heatmaps of ΔF/F traces of 13 brain regions (top) MFP activity in one complete male-female MFP recording session. Middle: HSMM-decoded neural sequence, which was derived based on the behavioral membership using the suboptimal state-behavior match. The latent sequence was also superimposed by the behavioral label color for improving visualization. Bottom: the annotated behavioral label. Three color-coded triangle symbols represent three different behavioral sequences. (B) The confusion matrix that shows the best match between latent neural states and behaviors. The WCP statistic is 0.918. (C) Inferred state-firing rate matrix. (D) Hierarchical clustering of inferred latent states revealed their relationship. (E) The newly inferred confusion matrix from truncating the duration of “*Other*” label. The WCP statistic is 0.905. (F) The newly inferred state relationship from truncating the duration of “*Other*” label. (G) The correspondence matrix between two sets of latent states inferred from two conditions showed clear correspondence. (H) The WCP population statistics from all MFP recording sessions (where the complete session was used for training only) according to three categories: male-male (n=16), male-female (n=14), and toy interactions (n=4). Red dots indicate statistical significance. (I) The WCP population statistics for a total of 31 testing epochs during male-male and male-female interactions. Red dots indicate statistical significance.

We further examined the inference outcome with respect to the presence of the “*Other*” category. Since the duration of the “*Other*” period was predominantly long, we reduced the duration of all annotated “*Other*” epochs such that the remaining duration matched the median duration of other behavioral states while keeping the transition boundary unchanged. Upon applying unsupervised HSMM learning on the truncated data, we further compared the inferred states with the annotated behavioral labels (**Figure 4E**) and examined the inter-state relationship (**Figure 4F**). The new WCP index was 0.905 and remained statistically significant (*P*<0.05). Importantly, a comparison between the latent states derived from these two conditions, with and without the whole “*Other*” period (**Figure 4B vs. 4E**), showed a clear correspondence (**Figure 4G**).

Among the total analyzed sessions (considered separately for male-male, male-female, and toy interactions), the majority of obtained WCP metrics were statistically significant for the non-truncated setting (**Figure 4H**). Some of insignificant sessions were likely due to the short duration of epochs, resulting in a lack of diversity in random shuffling. As a comparison, the WCP statistics from the testing epochs for the first type of analysis (where training and testing epochs were distinct) is shown in **Figure 4I**. Notably, the number of annotated behavior labels was much smaller in male-male interactions than in male-female interactions, which caused some performance discrepancy between the two conditions. In the special case where there was only one label (excluding the “*Other*” label), the WCP was reduced to 1.

### Post-hoc analysis validated HSMM-identified behaviorally relevant states

To understand potential micro-behavior states identified by the HSMM, we visually inspected the videos that correspond to different HSMM states and made several noteworthy observations in this male-female interaction session (see **Figure 5A** and **Movie S1** for the **Figure 4** male-female interaction example): (i) There were specific brain states during non-social interaction “*Other*” period signaling the progression of the overall male-female encounter. For instance, states 7 and 8 only occurred when the female intruder was first introduced; state 6 appeared during the non-interaction period after the initial introduction and before mounting started; and states 9 and 10 occurred during the non-interaction period after mounting started. (ii) Certain states corresponded to periods of distinct social behaviors. For example, states 4 and 5 represented “*Investigate*” whereas states 1-2-3-11 marked the behavioral sequence of “*Attempted mount*⤍*Mount*⤍*Thrust*⤍*Dismount*”’. (iii) Some neural states in the “*Other*” period corresponded to particular self-oriented behavior. For example, facial grooming was frequently observed during states 12 and 16.

**Figure 5.**
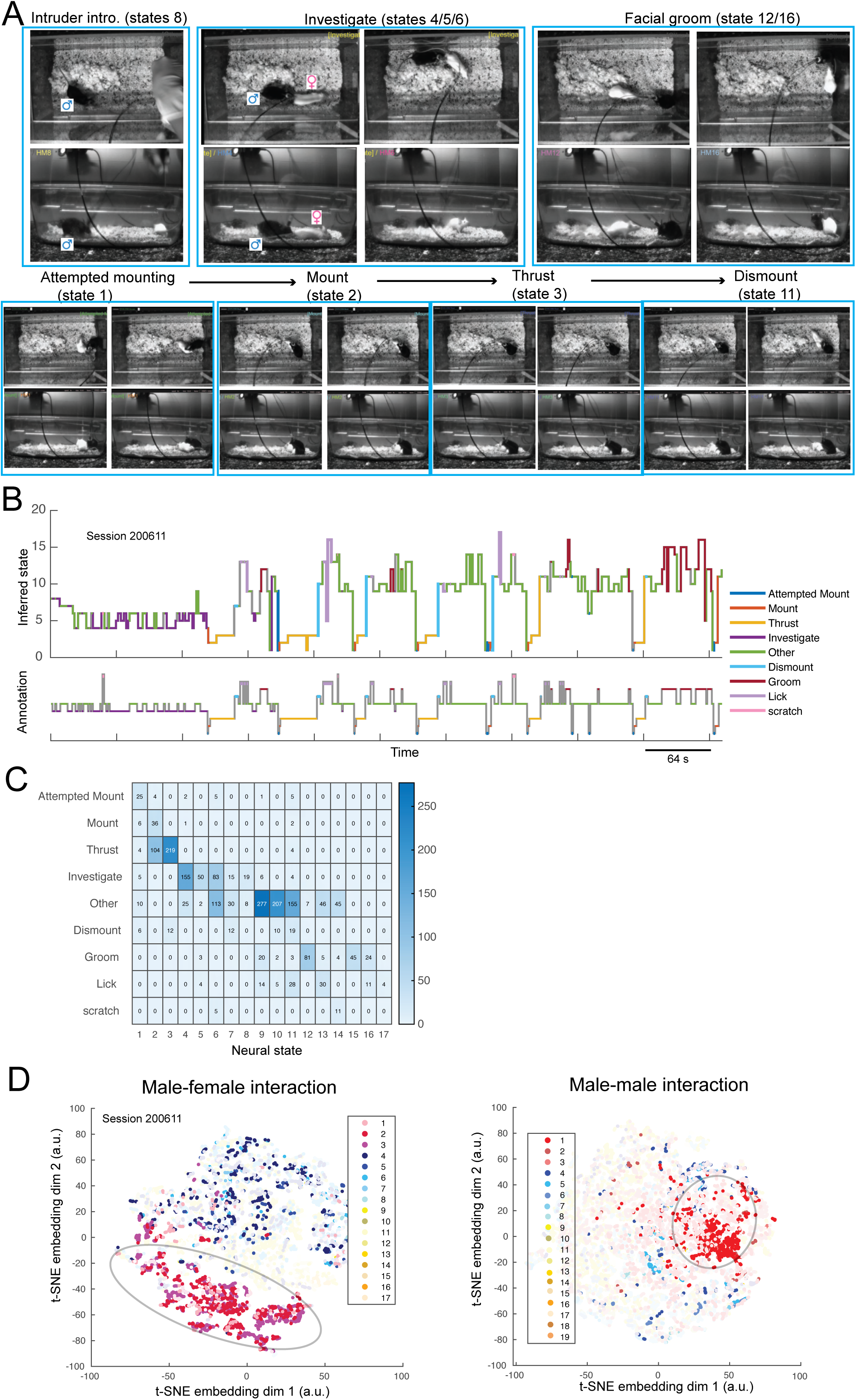
Post-hoc analysis validated HSMM-identified behaviorally relevant states. (A) Human inspection of video footage confirmed meaningful neural states from the HSMM inference. (B) HSMM-decoded neural sequence using the suboptimal state-behavior match based on the new annotations. The latent sequence was also superimposed by the behavioral label color for improving visualization. Bottom: the new annotated behavioral labels. (C) The new confusion matrix based on the new annotated behavioral labels. Compared to Fig. 4B, the state-behavior mapping was improved. The state-firing rate matrix remains unchanged from Fig. 4C. (D) Two representative male-female and male-male behavioral embeddings superimposed by the latent state identity with distinct color.

To better quantify the micro-behaviors associated with HSMM states, we conducted a more detailed frame-by-frame behavioral annotation (**Movie S1**), with four additional behavioral labels from the previous annotation: “*Dismount*”, “*Groom*”, “*Lick*”, and “*Scratch*”). Importantly, these annotations were performed blind to the HSMM state information. Next, we replotted the inferred neural sequence (same as **Figure 4A**) but with the new behavioral annotations (**Figure 5B**), and computed the confusion matrix (**Figure 5C**). Consistent with our initial impression, we found grooming, licking and scratching occur predominantly under specific and non-overlapping states. Compared to **Figure 4B**, the state-behavior mapping was improved in terms of grouping. As previously reported, it showed increased activity of BNSTp and MPN during mating (Guo et al. 2023). Interestingly, it revealed elevated activity in the PMv, a small hypothalamic region, during grooming and licking. Thus, the HSMM is capable of detecting neural signatures that generate specific social and non-social behaviors as well as specific motivational or emotional states that may not be manifested as discrete motor actions. This example session demonstrates the utility of the HSMM in objectively detecting behaviorally meaningful latent brain states and finding the relevant neural activity patterns.

To further reveal the relationship between brain and behavioral states, we employed a nonlinear dimensionality reduction technique (**Methods**) to compute a two-dimensional frame-by-frame behavioral embedding based on behavioral tracking variables for each session, and then colored the sample points according to the neural states. In general, the embeddings showed behaviorally meaningful clusters. For instance, in the case of male-female interaction (see the example in **Figure 4A**), the states 1-3 belonging to the “*Attempted mount, Mount, Thrust*” behaviors were mostly scattered in the bottom left corner, whereas states 4-6 that marked “*Investigate*” were mixed with “*Other*” states (left panel, **Figure 5D**); in the case of male-male interaction (see the example in **Figure S2**), states 1-3 mostly marked the “*Attack*” state, whereas states 4-7 representing the “Investigate” state were mixed with “*Other*” states (right panel, **Figure 5D**). In both cases, there were still varying degrees of overlap between aggressive/mating states and other micro-behavior states.

Finally, to assess the relative impact of the social network hub on the neural/behavioral states, we assumed that some brain regions were selectively inactivated and further applied maximum likelihood decoding to infer the most likely states (**Methods**). Taking the previous “*Attempted Mount*⤍*Mount/Thrust*” behavioral sequence as an example, we observed a predominant state transition (1⤍3, 1⤍2) or self-transition (1⤍1) among inferred neural sequences. In a simulated setting, we found that inactivating ABN did not change the mating behavior, whereas inactivating MBN could lead to the termination of mating behavior (**Figure S3**).

### HSMM analysis was robust with respect to data resampling and noise

An identifiability question naturally arises during the rank-invariant resampling procedure for MFP data: will the inference procedure produce a consistent state estimate regardless of the (nuisance) model parameters? To answer this question, we generated simulated ground truth data based on empirical statistics (such as the transition probability and mean amplitude from annotated behaviors) and ran extensive computer simulations using (i) various model hyperparameters and (ii) various additive white Gaussian noise distributions. For each condition, we generated a 5-by-13 state-firing template by computing the mean MFP activity in each behavioral category for each region. We then generated 1,000-5,000 data points according to a beh<u>s</u>aviorally relevant 5-by-5 state transition matrix (with five states representing {*Attempted Mount, Investigate, Mount, Other, Thrust*} during male-female interactions, but the five states can also generalize other behaviors). Finally, we performed sampling, resampling and potential noise addition to obtain the final simulated data (**Figure 6A**). The objective was to compute the consistency between the ground truth and estimated states and to quantify the variability in this consistency as a function of the model hyperparameter, sample size, and signal-to-noise ratio (SNR) (**Figure 6B**). Overall, the WCP index was rather stable across all conditions. In some examples (e.g., **Figure S4**), the number of inferred latent states was greater than the ground truth, but the neural states that co-represented true behaviors had similar neural patterns.

**Figure 6.**
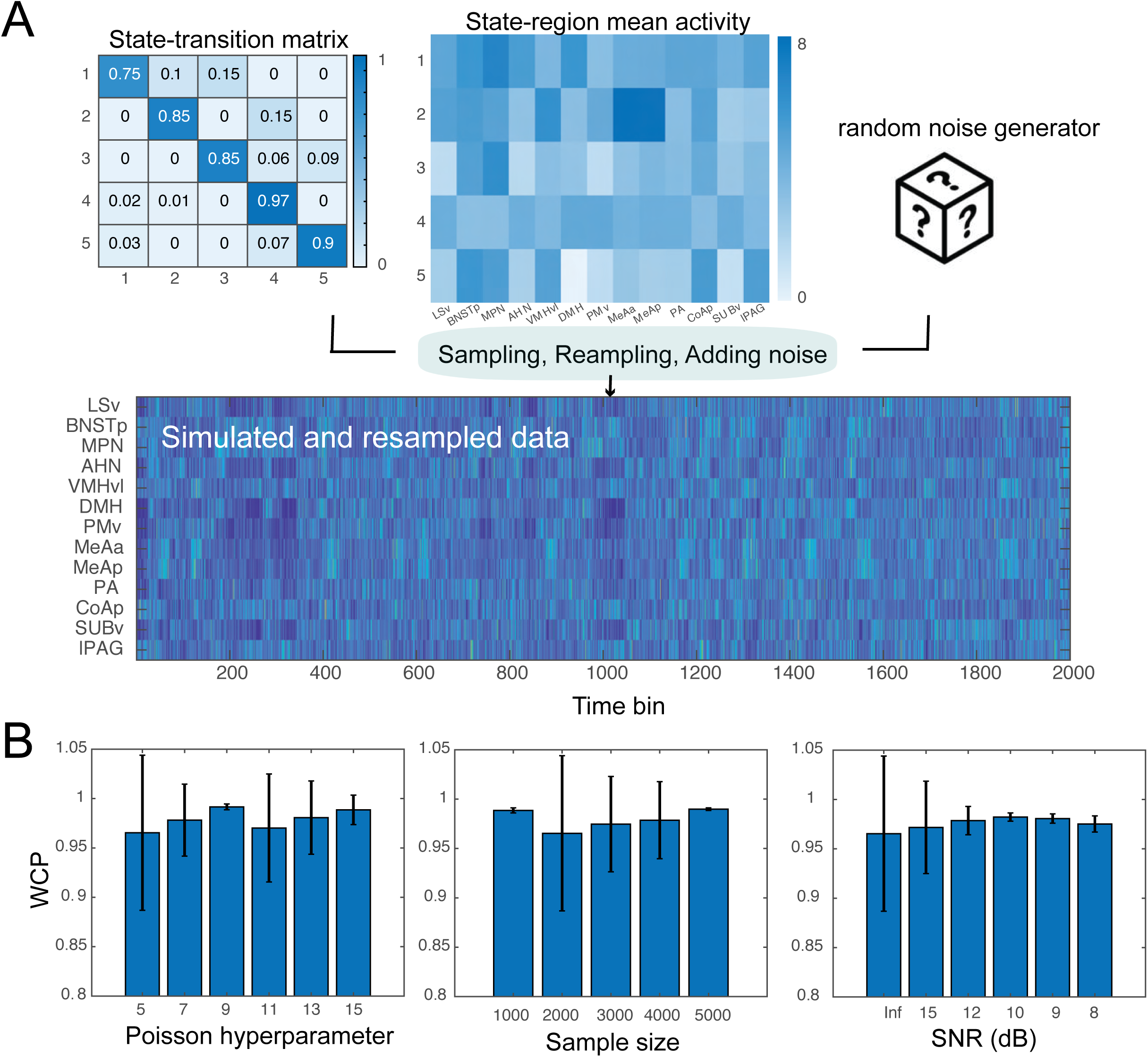
Latent state inference was robust with respect to resampling, sample size, and SNR. (A) Schematic of generating simulated data via sampling (from a user-defined state-transition matrix and a state-region mean activity template), rank-invariant resampling, and adding additive noise. One simulated 13-by-2000 data matrix is shown. (B) Comparison of WCP with respect to the resampling hyperparameter, sample size, and SNR. Error bar denotes SD over n=10 Monte Carlo experiments.

## DISCUSSION

Brain-wide multifiber photometry is an emerging technique that can identify and analyze functional networks across multiple brain regions, providing a versatile tool for studying large-scale cortical and subcortical brain dynamics during behavior (Gunnadyin et al., 2014; Sych et al., 2019; Formozov et al., 2023). The application of latent variable models for analyzing neural dynamics during complex animal behavior has become increasingly popular in neuroscience (Chen, 2015; Sani et al., 2021; Yang et al., 2021; Mazzucato, 2022). The LDS and HSMM approaches constitute two representative analysis paradigms. The latent neural states, whether discrete or continuous, can provide insight into the relationship between unobserved and observed variables. In our work, the LDS and HSMM approaches identified behaviorally relevant neural states, which were then linked to behavioral tracking data or annotated behavioral labels. In the literature, other related latent variable models have also been proposed. For instance, a combined generalized linear model-hidden Markov model (GLM-HMM) was used to identify animals’ latent cognitive states underlying complex decision-making tasks (Bolkan et al., 2022; Ashwood et al., 2022), or decode social competition behavior from mouse medial prefrontal cortex (mPFC) ensemble activity (Podilla-Coreano et al., 2022). Recently, a recurrent switching linear dynamical system (rSLDS) model was used to fit population activity of the mouse ventromedial hypothalamus (VMHvl) during social behavior to identify a low-dimensional neural state space (Nair et al., 2023). Our proposed analysis framework was conceptually in line with these studies but our effort focused on modeling gross multi-population activity instead of multiple single-unit activities.

In the LDS approach, we used a subspace system identification method (Sani et al., 2021) to identify behaviorally relevant latent states and further predicted neural activity and behavioral tracking data using a Kalman filter. While one-step-ahead prediction of neural activity was reliable, the behavioral prediction accuracy was less satisfactory, possibly due to multiple reasons: low-sampling rate nature of MFP signals, low quality of behavioral measures due to limited body-part tracking, and low spatial resolution of the tracking data. While this approach shows promising results in behavioral prediction, incorporating higher-order state transition dynamics into the LDS model or collecting MFP recordings from other behaviorally relevant brain regions may potentially improve the performance of behavioral prediction. The continuous latent variables also enable us to visualize neural subspaces and quantify the neural trajectories in a manner similar to that of the continuous line attractor model despite the difference in neural signal modality (Nair et al., 2023; Mountoufaris et al., 2023). Interestingly, our findings of rotated neural patterns around the onset of attack, mount, or thrust behaviors were shared between male-male and male-female interactions.

In the HSMM approach, we regarded MFP activity in each region as a proxy for a weighted sum of Poisson spiking activity in that region (Sych et al., 2019). However, it remains debatable whether photometry accurately reflects spiking-related and other changes in neuronal calcium (Legaria et al., 2022). A key difference between spiking and calcium activity is spatial and temporal resolution. Simultaneous in vivo electrophysiology and fiber photometry recordings in mouse dorsal striatum have shown that neural signals from spiking units are heterogeneous, whereas population calcium signals are homogeneous but display coordinating ramping activity preceding food approach and consumption (London et al., 2018).

The unsupervised HSMM approach can potentially identify micro-behavioral states (including during no-interaction periods) beyond those defined by human annotations. Therefore, such neural-based-behavior-state discovery could represent a key advantage of using the HSMM in addition to human annotation. Careful comparison of inferred state sequences and video footage revealed not only strong state-behavior correspondence but also new interesting behavioral labels (such as grooming or eating). Additionally, Bayesian nonparametric inference allows automatic identification of the number of such states. In practice, a specific annotated behavioral label may correspond to more than one neural state, reflecting some subtle differences in neural representations. Overall, the HSMM analysis provides a compelling proof-of-concept for exploratory analysis in neural-behavioral segmentation. While our current paper is more methodology-driven, a systematic investigation focusing on circuit and experimental questions will be presented in the near future.

The present study faces several limitations. First, our MFP recordings are restricted to 13 regions of hypothalamic-midbrain circuits; thus, spatial sampling of the SBN is limited. Indeed, it is likely that many other brain areas are also involved in innate mouse behaviors (see some reviews in Chen and Hong, 2018; Wei et al., 2021; Xu et al., 2021; Mei et al., 2023b). Future extension of MFP recordings to more brain regions can potentially improve behavioral decoding accuracy. Second, the fact that behavioral occupancy in freely behaving mice was not uniformly distributed creates challenges for statistical model identification and testing. In particular, the “*Other*” label was predominant in all recording sessions, yet the behavioral phenotypes associated with this umbrella category were complex and heterogeneous, making conclusive interpretations difficult. Third, behavioral ground truth supported by video tracking of two cameras was sometimes missing due to occlusion or limited camera view.

Overall, the present work contributes to a growing literature on neural-behavioral mapping in the mouse SBN (Falkner et al., 2020; Gao et al., 2023; Nair et al., 2023). Compared to electrophysiology, MFP recordings are capable of capturing population activity from a broad range of subcortical areas. Additionally, latent variable models allow us to characterize unobserved drivers of observed neural activity, such as top-down attention or common input. Identifying behaviorally relevant latent dynamics will improve our understanding of the link between hypothalamic-midbrain circuits and mouse social behaviors. Finally, our neural analysis framework may hold promise for the online detection of aggressive behavior and closed-loop experiments in socially interacting mice. Combining wireless MFP recording and closed-loop optogenetic stimulation may represent a future direction in dissecting large-scale social behavior circuits.

## METHODS

### Animals

Experimental mice for MFP recording were socially naïve, Esr1-2A-Cre male and female mice (10–24 weeks, Jackson stock no. 017911). After surgery, all test animals were single-housed. Intruders used were group-housed BALB/c males or group-housed C57BL/6 females (both 10-36 weeks, Charles River). Mice were housed at 18-23 °C with 40-60% humidity and maintained on a reversed 12-h light/dark cycle (dark cycle starts at 10 a.m.) with food and water available ad libitum. All experiments were performed in the dark cycle of the animals. All procedures were approved by the IACUC of New York University Grossman School of Medicine (NYUGSOM) in compliance with the NIH guidelines for the care and use of laboratory animals.

### Optical setup

The MFP arrays and recording system is as previously described (Gao et al. 2023). Briefly, the multi-fiber arrays were constructed using MT Ferrules (US Conec, No 12599) and 100 µm-core optic fibers (Doric Lens, NA0.37). The implanted arrays were connected with the MFP system through a custom-designed 19-fiber multi-fiber bundle (Doric Lenses, BFP(19)_100_110_1100-0.37_4m_FCM-19X). The MFP recording system delivers the blue excitation light (Thorlabs, M470F1, LEDD1B) to the end of the fiber bundle and collects and projects the emitted green light from the fiber bundle onto the CCD sensor of a camera (Basler, acA640-120um) via an achromatic doublet (Thorlabs, AC254-060-A-ML). The sampling rate of the camera was 25 frames per second.

### Stereotaxic surgery

Esr1-Cre mice were anesthetized with 1.5%-2% isoflurane and placed on a stereotaxic surgery platform (Kopf Instruments, Model 1900). 60-100 nl AAV2-CAG-FLEX-GCaMP6f (Vigene, custom prepared) or AAV1-CAG-FLEX-GCaMP6f (Addgene, 100835-AAV1, 3x dilution) was delivered unilaterally into each of the following targeted brain regions: LSv (AP 0.05, ML −0.65, DV −3.25); MPN (AP 0.00, ML −0.33, DV −4.90); BNSTpm (AP −0.30, ML −0.80, DV −3.60); AHN (AP −1.05, ML −0.55, DV −5.15); DMH (AP −1.75, ML −0.55, DV −5.2); VMHvl (AP −1.70, ML −0.75, DV −5.80); PMv (AP −2.40, ML −0.55, DV −5.70); MeAa (AP −1.10, ML 2.10, DV −4.90); MeApd (AP −1.60, ML 2.10, DV −4.92); PA (AP −2.35, ML 2.20, DV −4.92); CoApm (AP −2.85, ML 2.90, DV −5.20); SUBv (AP −3.35, ML 2.60, DV −4.60); lPAG (AP −4.90, ML −0.45, DV −2.40). All regions included in the final analyses had correct virus expression and fiber tip position based on histology.

For each animal, two custom-made multi-fiber arrays were implanted, one designed to target seven medial regions on the left hemisphere and the other to target five lateral regions on the right hemisphere. A custom-made optic-fiber assembly targeting lPAG (Thorlabs, CFX126-10) was also implanted in the same animal. All optic fibers are targeted ∼250 μm above the injection sites and secured using dental cement (C&B Metab ond, S380). lPAG fiber was implanted at (AP −5.20, ML −0.45, DV −2.00) after tilting the head 8 degrees down rostrally to avoid collision with the other two arrays. Lastly, a 3D-printed plastic ring for head fixation was cemented on the skull (Osborne and Dudman, 2014).

### Animal behavioral tracking and annotation

Behaviors were recorded under dim room light via two cameras from top and side views (Basler, acA640-100gm) using StreamPix 5 (Norpix), which also coordinated the MFP camera in synchrony. Animal positions were tracked using DeepLabCut software (v.2.0.6) (Mathis et al., 2018). The tracking data consisted of 2D positional information from seven points on the bodies of two mice. These seven points are the left ear, right ear, nose, center, lateral left, lateral right, and tail base. Then, the convex hull mentioned below was drawn on a 2D plane using the positions of several points on the mouse’s body. A total of 29 behavioral features (**Table S2**) were extracted from computerized video analysis using custom MATLAB routines. Furthermore, manual behavioral annotations were made by trained experimenters on a frame-by-frame basis, where human observers performing annotations were blind to experimental conditions and recordings. Annotated behaviors were categorized as follows:

‘*Investigation*’, the resident mouse made nose contact with either the facial or anogenital region of the intruder mouse or the whole body of the toy mouse. “Investigation” also consists of “*Male Investigate*” or “*Female Investigate*”.

‘*Attack*’, a suite of actions initiated by the resident toward the male intruder, which included lunges, bites, tumbling and fast locomotion episodes between such behaviors.

‘*Attempted Mount*’, began when the resident male charged toward the rear end of the female body, rose and grasped the female’s flank with his forelimb, and ended by aligning his body with the female’s and assuming the on-top posture.

*’Mount’*, the male grasped the female’s body tightly with his forelegs and made rapid shallow pelvic thrusts.

‘*Thrust*’, the male made deep rhythmic movement of pelvis presumably with penile insertion into the vagina.

‘*Ejaculation*’, the male froze at the end of an intromission event while continuously clutching onto the female and then slumping to the side of the female. Ejaculation occurred only once in a female session, signaling the end of sexual behaviors. It was always confirmed by the presence of a vaginal plug after the recording.

*’Other’*, other remaining time points, including many during which the two animals were separate from each other and not closely interacting.

Specifically, the behavioral labels {*Attack, Investigate*} were usually associated with male intruders, whereas {*Investigate, Attempted Mount, Mount, Thrust, Ejaculate*} were usually associated with female intruders. In total, there were 7 annotated behavioral events across all analyzed sessions (**Table S1**), with a varying number of behavioral labels in each session. We computed the mean duration of each labeled behavior, and between-behavior transition probability matrix. This information was used to determine the bin size and generate computer simulation data.

### Multi-fiber photometry (MFP) recording

The recording started three weeks after the virus injection. For each recording session, the head-implanted MT ferrules were connected to the matching connectors at the end of the custom optic bundle (Doric lens, BFP(19)_100/110/1100-0.37_4m_SMA-19x). A drop of liquid composite (Henry Schein, 7262597) was applied to the outer part of the junction and cured with blue LED curing light (Amazon) to stabilize the connection. The baseline signal was checked in the absence of the intruder for at least two days to ensure that the signal reached a stable level (<10% difference across days for all regions). On the recording day, after 10 minutes of the baseline period, a sexually receptive female mouse was introduced until the recording male achieved ejaculation or after 60 minutes. Then, 5 minutes after removing the female, a group-housed non-aggressive BALB/c was introduced for 10 minutes. For some recording sessions, 5 minutes after removing the male, a novel object (15 mL plastic tube) was introduced for 10 minutes. Each animal was recorded 2-4 times, with at least three days in between. The order of male and female presentations was counterbalanced across sessions. In total, we analyzed 22 sessions from 5 animals (**Table S1**).

### Preprocessing and resampling

From MFP recordings, regions of interest (ROIs) for selected channels were drawn on the grayscale image of the optic fiber bundle, and the average pixel intensity for each ROI was calculated as a readout of the raw Ca^2+^ signal (F) for the region. We then computed the change in fluorescence (ΔF/F) as (F_raw_-F_baselline_)/F_baseline_, where F_baseline_ denotes the instantaneous baseline. All follow-up analyses were derived from ΔF/F, followed by Z-scoring within the same region in the same session. In calcium imaging, spike deconvolution methods are often applied to infer the proxy of spiking activity of single neurons. While MFP recordings do not have cellular resolution, the same principle still holds. From the individual Z-scored ΔF/F traces, we further employed a peak detection algorithm to detect the discrete spiking activity as filtered point process (FPP) events that produced a good approximation to deconvolved calcium fluorescence (Tu et al 2019). Briefly, we defined a peak as any time point with a value greater than both its preceding and following neighbors and set to 0 any time points that were not peaks. We then convolved an exponential filter across the timeseries to capture ramping spiking activity. We further assumed that the fiber photometry signal is a proxy for the underlying population activity (Sych et al 2019, Willmore et al 2022). As a result, the recovered nonnegative signal should be a *weighted* sum of independent Poisson random variables, which is approximately Poisson (note that the unweighted sum of independent Poisson variables is exactly Poisson distributed).

We next binned the data using either 320 or 280 ms bin size such that each bin still contained at most one transition between the annotated behavior. We further applied a rank-invariant resampling procedure to transform a univariate continuous measure into a univariate discrete counting measure while preserving the rank order (i.e., if x_1_>x_2_, X_1_>X_2_, where X_1_ and X_2_ denote the resampled data points from original data x_1_ and x_2_, respectively). A schematic illustration of the resampling procedure is shown in **Figure 1F**. As a hyperparameter, a larger Poisson mean in rank-invariant resampling implies a larger Poisson variance. However, the result of state inference was robust with respect to a wide range of the Poisson mean statistics. In most of our experiments, we resampled data points from each brain region independently from a Poisson distribution with a mean parameter of 5.

### Hidden Markov and semi-Markov models

To characterize the neural dynamics that drive an animal’s behavior, we modeled the abstract neural state as a Markovian or semi-Markovian process with a switching mechanism. In a simple setting, we assumed that the latent state process followed a first-order discrete-state Markov chain {𝑆*_t_*} ∈{1,2,···,*N_s_*}, and that the observations of neural activity at discrete time index *t* followed a conditional probability distribution (conditioned on the latent state 𝑆*_t_*). If the observations are discrete counting measures, a Poisson or gamma distribution is a natural choice, which can be parametrized by 𝜦. For a set of time points *t*=1,2,··,*T,* the joint probability distribution of observed and latent variables is given by

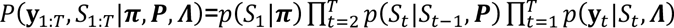

where 𝑷 = {𝑃*_ij_*} denotes an *N_s_*-by-*N_s_* state-transition matrix, with 𝑃*_ij_* representing the transition probability from state *i* to *j*; 𝝅 = {𝜋*_i_*} denotes a probability vector for the initial state *S*_1_; *y_c,t_* denotes the observation from the *c*-th brain region at time *t*. If the conditional distribution 𝑝(𝐲*_t_*|𝑆*_t_*) is a Poisson distribution, we can write 𝜦 as a compact form as a *C*-by-*N_s_* matrix. We varied the temporal bin size between 120 ms and 400 ms. To allow computationally efficient inference, we assumed conjugate priors for both parameter 𝝅 and 𝜦. Following a hierarchical Bayesian formulation of the HMM, HDP-HMM, we derived Markovian chain Monte Carlo (MCMC) sampling-based inference algorithms to estimate latent states and model parameters { 𝝅, 𝑷, 𝜦 } (Linderman et al., 2016).

To accommodate a history dependence on state switching, we extended the hidden Markov model (HMM) to a hidden semi-Markovian model (HSMM), which assumed that *P_ij_* depends on the sojourn duration in state 𝑖 (Linderman et al., 2016). One possibility is to introduce a sticky HDP-HMM (Fox et al., 2011) by adding a global hyperparameter 𝜅 to encourage the probability of self transitions. However, since 𝜅 is applied to all latent states, the HMM may still struggle to learn latent states with substantially different durations; further, the distribution of state durations remains geometrically distributed (Johnson and Willsky, 2013). Here, we introduced an explicit-duration semi-Markov modeling for each latent state and considered latent “super-states.” This was implemented by assuming that the sojourn duration in state 𝑖, denoted by *p*(*d_t_*|*S_t_ =* 𝑖), followed a parametric distribution form:

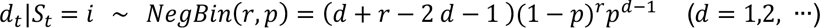

where 𝑁𝑒𝑔𝐵𝑖𝑛(𝑟, 𝑝) denotes a negative binomial distribution, which reduces to the geometric distribution when 𝑟 = 1 (i.e., under the Markovian assumption). We used an “embedding trick” for the negative binomial distribution (Johnson and Willsky, 2013; Chen et al., 2016). A close examination of sojourn duration distributions in HSMM allowed us to validate the semi-Markovian dynamics.

### Performance assessment

To assess the performance of the HSMM, we split the MFP data in each recording session between training and testing sets by accounting for multiple factors: temporal continuity as well as occurring occupancy and frequency of behavioral actions. The ratio of training to testing data points varied from individual sessions. In the post-hoc testing, we first matched model hidden states to behavioral labels. However, the former always outnumbered the latter, and many classic approaches to the assignment problem, such as the Hungarian algorithm, are designed to match two classes of equal size and may leave items unassigned if the sizes of the classes differ. In order to assign every hidden state to a behavioral label, we thus first computed the count of overlapping bins between each behavior and state. We then performed two rounds of sorting. We first matched each hidden state to the one behavioral state with which it maximally overlapped; finally, for each behavior, all states matched to that behavior were ordered by the strength of their overlap. Further, we summarized the correspondence between the inferred neural states (x-label) and behavioral labels (y-label) with a confusion matrix. Each column specified how each neural state represented distinct annotated behavioral actions, whereas each row specified how each action was represented by different neural states. A one-to-one correspondence implies no ambiguity, whereas column “impurity” indicates state non-specificity. In contrast, row “impurity” implies behavioral variability that calls for multiple neural states. For each session, depending on the occupancy of each annotated behavior, we computed a weighted column purity metric:

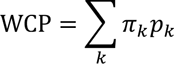

where 𝜋*_k_* denotes the relative behavioral occupancy, and 𝑝*_k_* denotes the column-normalized probability in the confusion matrix associated with the *k*-th behavior label among all non-”*Other*” labels. For WCP statistics derived from all possible paired training-testing epochs, we computed Pearson’s correlation.

To assess statistical significance of the derived WCP metric, we generated a randomly shuffled testing data matrix by using both row and column shuffle operations. In each row, we randomly shuffled and destroyed the temporal structure. In each column, we randomly permuted the indices to shuffle the brain region. We used the trained HSMM on the shuffled data to compute the WCP metric at the chance level. This procedure was repeated 1,000 times to generate a Monte Carlo *P*-value. We reported the *P*-values of all test epochs among all recording sessions.

### Quantification of latent state dissimilarity and hierarchical clustering

Based on the *C*-by-*N_s_* state-wise Poisson firing rate matrix 𝜦 inferred from the HDP-HMM, we defined a divergence metric, *D*(*S_i_, S_j_*), to characterize the dissimilarity between two latent states *S_i_* and *S_j_* (with associated firing rate vector 𝜆*_i_* = [𝜆_1,*i*_, …, 𝜆*_C,i_*] and 𝜆_1,*j*;_ = [𝜆_1,*j*_, …, 𝜆_*C,j*_])

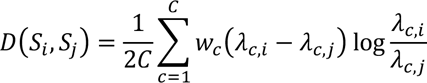

which is a weighted Kullback-Leibler (KL) divergence between two univariate Poisson distributions with distinct firing rates (Liu et al., 2018). Furthermore, we computed an *N_s_*-by-*N_s_* divergence matrix whose (*i,j*)-th entry defines the divergence between two states *S_i_* and *S_j_*. The diagonal elements of the matrix are all zeros.

Hierarchical clustering groups data over a variety of scales by creating a cluster tree or dendrogram, a multilevel hierarchy in which clusters at one level are joined to form clusters at the next level. Using the KL divergence metric, we applied a hierarchical clustering algorithm (MATLAB function: “clusterdata.m”) to define the similarity between the inferred latent states. The clustering procedure consisted of three steps: (i) define the similarity or dissimilarity between every pair of data points or strings in the data set; (ii) group the data or strings into a binary, hierarchical cluster tree; (iii) determine where to cut the hierarchical tree into clusters.

### Maximum likelihood inference for latent states

Based on the inferred state-firing rate matrix and resampled (or manipulated) Poisson activity from *C* brain regions, assuming conditional independence between brain areas, we applied the maximum likelihood method to infer the most likely state 𝑆_ML_

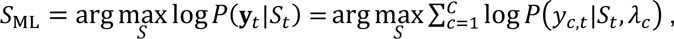

where 𝑃(𝑦𝑐,𝑡|𝑆_𝑡_, 𝜆_𝑆_) denotes the Poisson likelihood function with associated rate parameters 𝜦 = {𝜆_𝑐_}.

### Linear dynamical system (LDS) and preferential subspace system identification (PSID)

Following the neural-behavior joint modeling framework (Sani et al., 2021), let 𝐲*_t_* denote the neural activity and 𝐳*_t_* denote the behavioral activity. We assume that 𝐱_*t*_ is a continuous-valued latent state variable, where 𝐱*_t_* = [𝐱_1,*t*_, 𝐱_2,*t*_] is decomposed into two components such that *dim*(𝐱*_t_*)=*dim*(𝐱_1,*t*_)+*dim*(𝐱_2,*t*_), where 𝐱_1,*t*_ is assumed to be behaviorally relevant, whereas 𝐱_2,*t*_ is not. We further assume that the dynamics of variables {𝐱*_t_*, 𝐲*_t_*, 𝐳<u>_t_</u>} are characterized by a linear dynamical system:

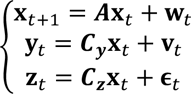

where {𝐰*_t_*, 𝐯*_t_*, 𝛜*_t_*} denote the Gaussian noise variables. Projecting future behavioral activity 𝐳*_t_* (denoted by *Z_f_*) onto past neural activity 𝐲*_t_* (denoted by *Y_p_*) enabled us to assess *Y*⤍*Z* predictability.

In our analysis, we assigned the 13-dimensional Z-scored ΔF/F to vector 𝐲*_t_*, and combined 7 features to form vector 𝐳*_t_*:

“*subj1_median_last_longest_hull”:*

Median of longest distance between any two points in the convex hull of the resident mouse in the last 166 ms.

*“subj1_median_last_centroid_mvmt”:*

Median centroid movement of the resident mouse in last 500 ms

*“subj1_median_last_tail_base_mvmt”:*

Median tail base movement of the resident mouse in the last 500 ms

*“both_mean_last_all_bp_mvmt”:*

Mean movement of all body parts of both subjects in last 166 ms

*“both_prctile_rank_mean_last_sum_bp_dist”:*

Percentile rank of mean (over last 500 ms) of sum of distances between respective body parts of the two subjects

*“subj2_shortest_last_median_bp_dist”:*

Shortest value (within sliding 500 ms window) for median pairwise body part distance for subject 2

*“subj1_prctile_rank_last_sum_centroid_mvmt”:*

Percentile rank of the sum of centroid movements of subject 1 over the last 500 ms

The model parameters could be efficiently estimated using a two-stage preferential subspace system identification (PSID) algorithm (https://github.com/ShanechiLab/PSID), which is a method that models neural activity while dissociating and prioritizing its behaviorally relevant dynamics. During learning, PSID first extracts the latent states directly using the neural activity and behavior training data, and then identifies the model parameters using the extracted latent states. In decoding, based on the LDS formulation, we used a recursive Kalman filter to predict 𝐲*_t_*_+1_ (one-step ahead self-prediction) and 𝐳_*t*+1_(one-step ahead behavioral prediction) based on historical neural activities 𝐲_1_to 𝐲*_t_*. The hyperparameters of the PSID algorithm were selected by grid search using cross validation (**Figure S1**). We also observed that using more behavioral tracking data did not qualitatively change the main findings.

To visualize the low-dimensional neural trajectory 𝐱*_t_*, we adopted a similar strategy as (Yu et al., 2009) and conducted a singular value decomposition (SVD) on the matrix 𝑪_𝒛_ = 𝑼𝑺𝑽′. We then transformed the latent variable 𝐱_*t*_ to 𝐱̃_*t*_ according to 𝐱̃_*t*_ = 𝑼𝐱_*t*_, where the columns in 𝑼 were sorted by their associated singular values (diagonal entries in matrix ***S***) in decreasing order. For visualization, we plotted the first two columns of 𝐱̃_*t*_ against each other (**Figure 2G**).

Given the dynamic transition matrix ***A***, we could compute the eigenvalues {𝜆 } (complex-valued), which reflect the rate at which activity along each latent dimension decays to zero following external input and which can be converted to a time constant for the *i*-th dimension (Nair et al., 2023): 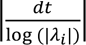 (where *dt* denotes the discrete-time bin size). The smaller the eigenvalue in absolute value (but within the unit cycle), the smaller is the time constant.

### Online prediction of attack behavior

In the LDS approach, we employed two nonparametric classifiers, random forest (RF) and gradient boosting machine (GBM), to discriminate “*Attack*” and non-Attack behaviors. We used the MATLAB “TreeBagger” and “fitcensemble” functions in the Statistics and Machine Learning Toolbox, with the “GradientBoosting” method specified for GBM. To optimize the performance of the GBM algorithm, the sample weights for “*Attack*” behaviors were increased to the quotient of the number of non-attack samples over attack samples, while the sample weights for non-attack behaviors were set to 1. Based on annotated labels, we first trained each classifier on selected sessions and tested the prediction performance on a new recording session based on a leave-one-session-out strategy. In training, the input features were either purely neural observations (N=13) or joint neural-behavior (N=20) observations. In testing, we used the Kalman filter to produce one-step prediction of neural observations or neural-behavior observations as the test input.

To assess the performance, we computed the standard metrics such as precision 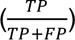, recall 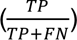, F1-score (i.e., harmonic mean of precision and recall), area under the ROC curve (AUC), and false alarm rate (i.e., number of false positives normalized by male-male interaction duration). While AUC was computed based on the per-frame prediction scores generated by the classifiers, precision and false alarm rate were computed accounting for possible boundary conditions by downsampling 25 frames to compare the prediction and labels. With this criterion, predictions of “*Attack”* that deviated from annotations by less than one second were not considered as false alarms.

### Principal component analysis (PCA) and contrastive PCA (cPCA)

Principal component analysis (PCA) was applied to analyze the Z-scored ΔF/F traces from 13 brain regions during “*Attack*” behavior in MFP recording sessions. Briefly, let us assume 𝐂_1_ and 𝐂_2_ are covariance matrices calculated from neural activity during a target and a background period, respectively. The cPCA algorithm (Abid et al., 2018) seeks to find a direction 𝐯 (also called contrastive principal component) to maximizes the function 𝐿(𝐯) = 𝐯*^T^*(𝐂_1_ − 𝛼𝐂_2_)𝐯, where 𝛼 is a contrastive parameter representing the trade-off between having the high target variance and the low background variance.

### Unsupervised behavioral embedding

Based on 29-dimensional behavioral features (**Table S2**), we applied the t-SNE (stochastic neighborhood embedding) algorithm (MATLAB “tsne.m” function) to extract two-dimensional (2D) behavioral embedding features. To visualize the relationship between latent neural states and behavioral states, we superimposed the discrete state ID (from the HSMM) with distinct color.

### Quantification and statistical analysis

All statistical analyses were performed using MATLAB or Python. Error bars are shown as means ± s.e.m. or means ± SD. We used two-sided Wilcoxon signed-rank or rank-sum tests for all paired and unpaired statistical tests, respectively. These tests are nonparametric and do not assume a specific distribution for the data.

## Data and code availability

The data used in our current analysis are publicly available https://zenodo.org/records/8128564. The custom software written in Python or MATLAB are also publicly available (https://github.com/YiboChen2020/MFPBehavioralAnalysis), and any additional request is available upon request from Dr. Zhe Sage Chen.

## ACKNOWLEDGMENTS

We thank the members of Lin lab for critical feedback on this manuscript. The work was partially supported by grants MH118928 (Z.S.C.), DA056394 (Z.S.C.), NS123928 (Z.S.C.), MH132642 (Z.S.C.), HD092596 (D.L.), MH101377 (D.L.), MH124927 (D.L.), and NS107616 (D.L.) from the National Institutes of Health.

## AUTHOR CONTRIBUTIONS

Z.S.C. and D.L. conceived and supervised experiments, and interpreted the data. Y.C. and J.C. performed simulations, analyzed and interpreted the data. B.D. assisted with behavioral video analyses. D.L. provided the experimental data. Z.S.C. and D.L. acquired funding. Z.S.C. wrote the paper with additional editing from D.L. and J.C.

