## Supplementary material for "Identifying behavioral links to neural dynamics of multifiber photometry recordings in a mouse social behavior network": Supp Figures and tables

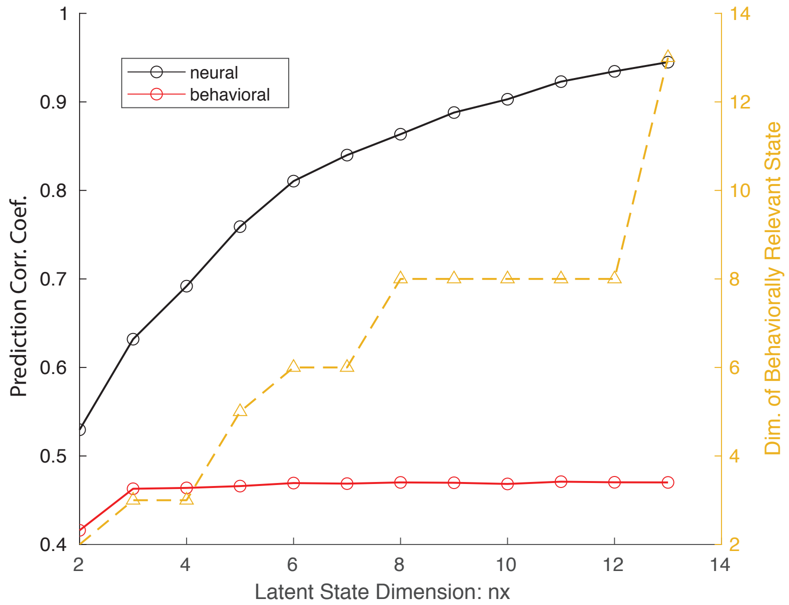


**Figure S1. Impact of hyperparameter choice on neural and behavioral prediction performance (average across 13 neural variables and 7 behavioral variables, respectively).** Generally, increasing the latent stat dimension nx=$dim(\mathbf{x}_t) \leq 13$ and behaviorally relevant state dimension $dim(\mathbf{x}_{1,t}) \leq dim(\mathbf{x}_t)$ gradually improved the prediction accuracy of neural observations, but saturated quickly in behavioral prediction.

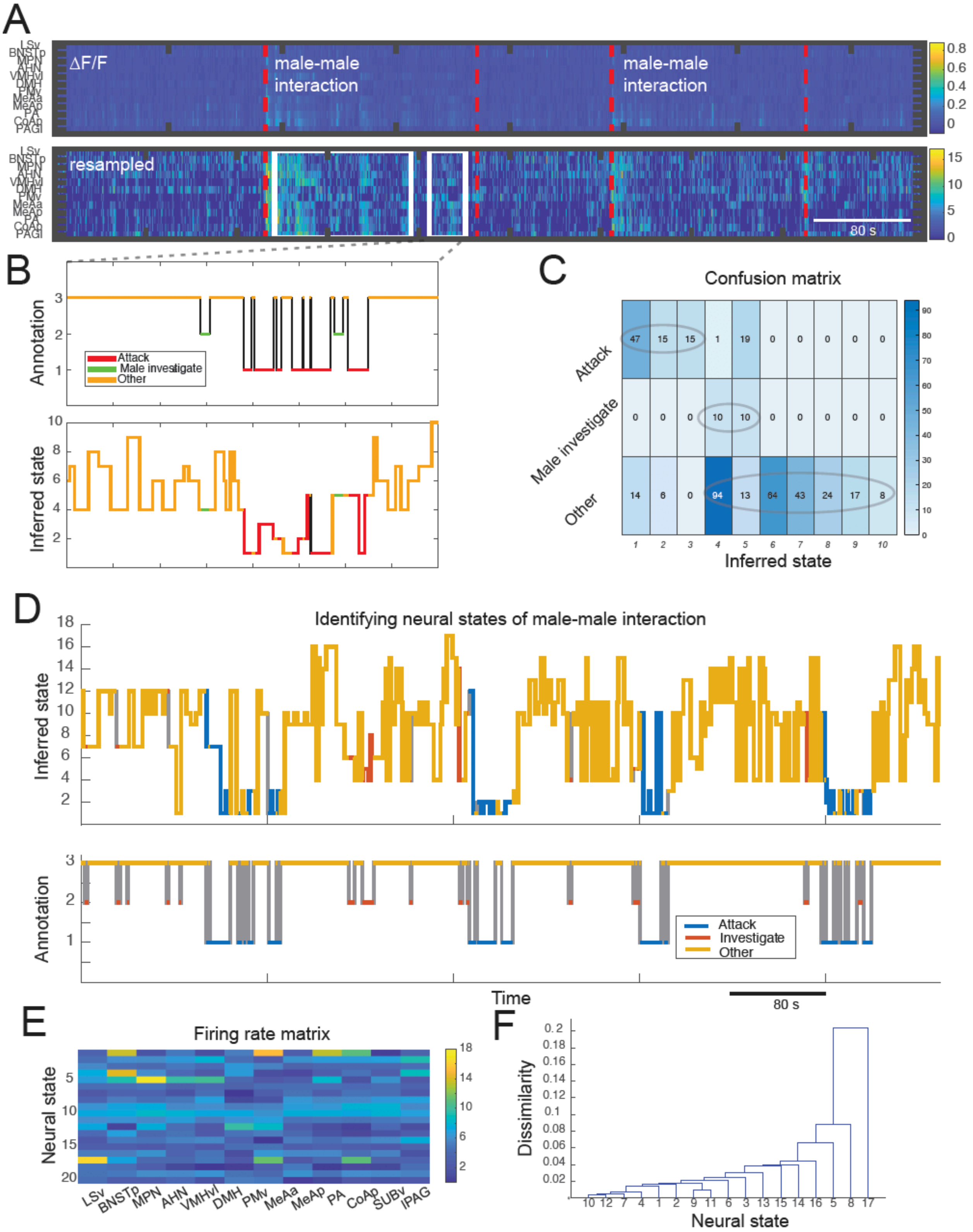


**Figure S2. HSMM estimation results from a representative male-male interaction training epoch and tests on an unseen male-male interaction epoch.**

(A) Heatmaps of ΔF/F traces of 13 brain regions (top) in one recording session. Vertical red dashed line indicates the onset of male-male interaction by introducing a male intruder. Two white boxes represent the selected training and testing epochs that share common annotated behaviors.
(B) Comparison of annotated behavioral sequence (top) and inferred state sequence (bottom) from the selected test epoch in panel A.
(C) Confusion matrix derived from the selected test epoch example in panel A.
(D) Top: HSMM-decoded neural sequence for identifying male-male interactions based on the whole MFP recording. The latent sequence was derived based on the behavioral membership using the suboptimal state-behavior match. The latent sequence was also superimposed by the behavioral label color for improving visualization. Bottom: the annotated behavioral label.
(E) Inferred state-firing rate matrix.
(F) Hierarchical clustering of inferred latent states revealed their relationship.

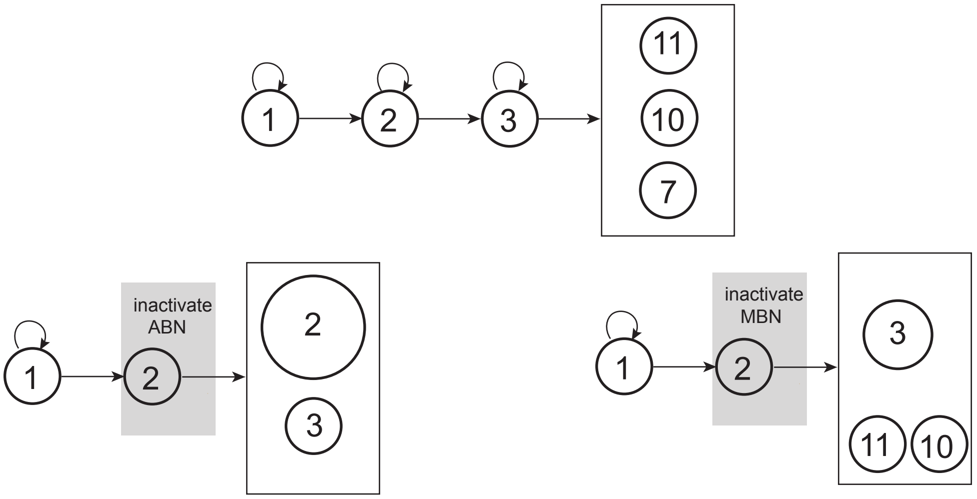


**Figure S3. Graphical illustration of neural state transition and the impact of inactivation of two network hubs: aggression-biased network (ABN) and mating-biased network (MBN).** Top: normal state transition 1→2→3 represents "*Attempted Mount→Mount→Thrust*", state 11 represents "Dismount", state 10 represents "*Other*", and state 7 represents "*Investigate*". Bottom left: inactivation of ABN does not affect the mating behavior sequence. Bottom right: inactivation of MBN increases the chance to transition out of the mating behavior.

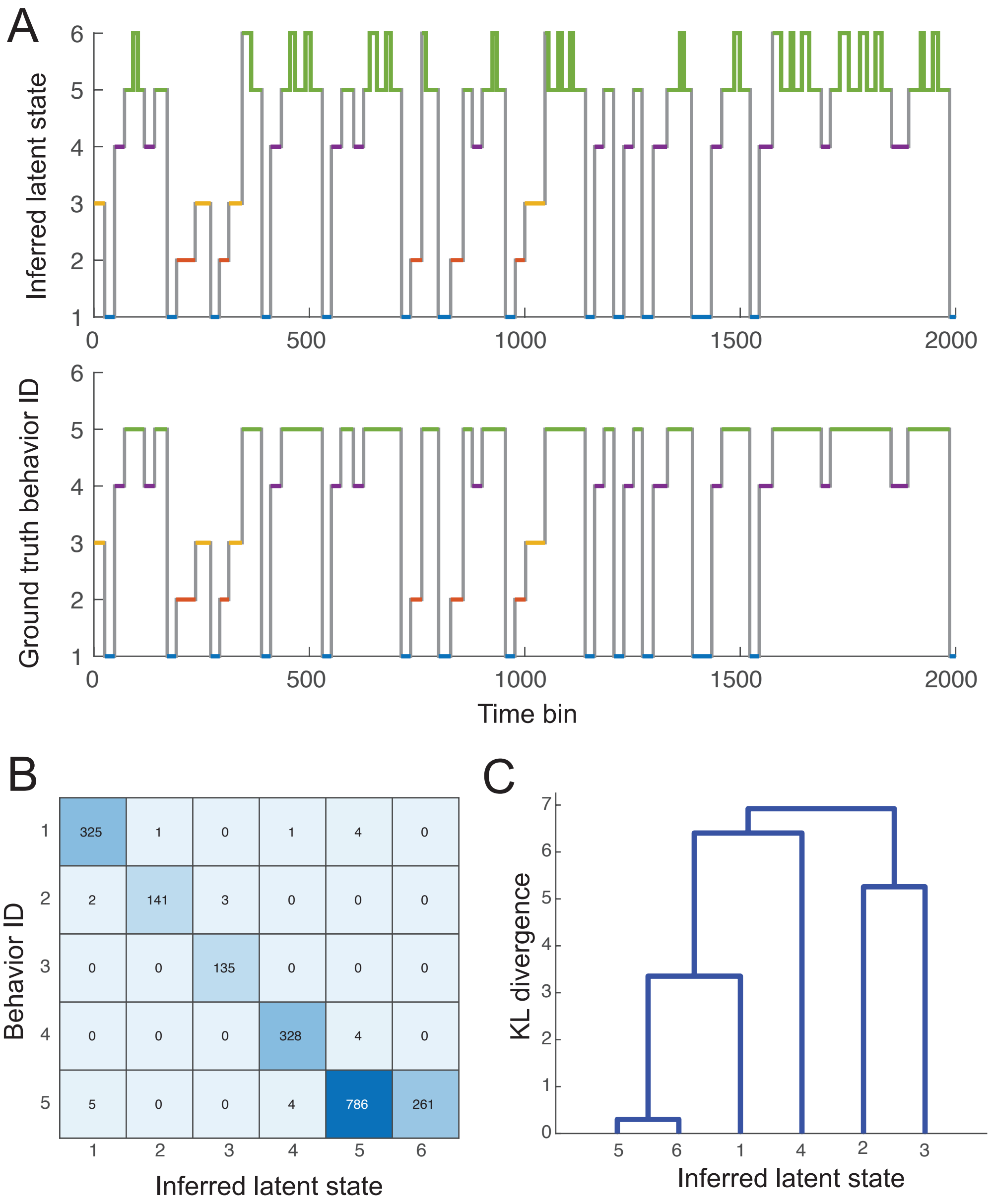


**Figure S4. Illustration of one computer simulation result.**

(A) The inferred latent state sequence (after greedy matching) versus ground truth behavior ID. Each behavior ID is coded with different color. The same color also was superimposed on the top panel.

(B) Confusion matrix between inferred latent states and behavior ID shows nearly one-to-one correspondence.

(C) Hierarchical clustering between states according to their dissimilarity. States 5 and 6 are similar as they both matched the same behavior (ID #5).

**Table S1. A summary of 22 MFP recording sessions used in the current study.**

| Animal | Session ID | Duration (s) | Number of annotated behavioral labels | Behavioral tracking |
|---|---|---|---|---|
| #1 | 190912 | 2882 | 6, No *Attack* | No |
| #1 | 190922 | 2900 | All 7 | Yes |
| #1 | 190925 | 3181 | 6, No *Male Investigate* | Yes |
| #1 | 191010 | 3883 | All 7 | Yes |
| #6 | 200311 | 4058 | 4, No *Thrust, Ejaculate, Male Investigate* | Yes |
| #6 | 200312 | 4001 | 3, No *Attempted Mount, Mount, Thrust, Ejaculate* | Yes |
| #6 | 200313 | 4930 | 2, *No Female Investigate, Attempted Mount, Thrust, Ejaculate, Male Investigate* | Yes |
| #6 | 200314 | 5363 | All 7 | Yes |
| #6 | 200315 | 4567 | 6, No *Ejaculate* | Yes |
| #7 | 200527 | 4312 | 6, No *Attack* | No |
| #7 | 200530 | 2588 | 4, No *Ejaculate, Male Investigate, Attack* | No |
| #7 | 200617 | 4201 | 5, No *Thrust, Ejaculate* | No |
| #8 | 200527 | 3901 | No *Attack* | No |
| #8 | 200611 | 4951 | 5, No *Ejaculate, Attack* | No |
| #8 | 200615 | 4803 | 6, No *Ejaculate* | Yes |
| #8 | 200617 | 3627 | 3, No *Attempted Mount, Mount, Thrust, Ejaculate* | Yes |
| #12 | 200801 | 6243 | All 7 | Yes |
| #12 | 200803 | 3601 | 6, No *Ejaculate* | Yes |
| #12 | 200827 | 3930 | 1, No *Female Investigate, Attempted Mount, Mount, Thrust, Ejaculate, Male Investigate* | Yes |
| #17 | 200708 | 4520 | 5, No *Thrust, Ejaculate* | Yes |
| #17 | 200709 | 5232 | 6, No *Male Investigate* | Yes |
| #17 | 200719 | 3618 | 6, No *Male Investigate* | Yes |
| | | Mean ± SD: 4150±891 | | |

**Table S2. A summary of animals' behavioral tracking features.**

| Feature No | Behavioral features of animals S1 (resident) and S2 (intruder) |
|---|---|
| 1 | Distance from S1 nose to S2 tail |
| 2 | Distance from nose to nose |
| 3 | Distance from S1 nose to S2 lateral left |
| 4 | Distance from S1 nose to S2 lateral right |
| 5 | Distance from S1 lateral left to S2 nose |
| 6 | Distance from S1 lateral right to S |
| 7 | S1 ear to ear distance |
| 8 | Sum of S2 width in last 500 ms |
| 9 | Sum of S1 width in last 500 ms |
| 10 | Median of longest distances on S1 convex hull in last 166 ms |
| 11 | Deviation from mean shortest distance on S2 convex hull in last 166 ms |
| 12 | Deviation from mean shortest distance on S2 convex hull in last 500 ms |
| 13 | Percentile rank of mean shortest distance on S2 convex hull in last 500 ms |
| 14 | S1 centroid X position |
| 15 | S1 centroid Y position |
| 16 | S2 centroid X position |
| 17 | S2 centroid Y position |
| 18 | Percentile rank of distance between S1 and S2 centroids |
| 19 | Percentile rank of sum of distances between all respective body parts of S1 and S2 |
| 20 | Median S1 centroid movement speed in last 500 ms |
| 21 | Median S1 tail base movement speed in last 500 ms |
| 22 | Sum of shortest pairwise distance between S2 body parts in last 166 ms |
| 23 | Mean of shortest pairwise distance between S2 body parts in last 166 ms |
| 24 | Mean movement speed of all body parts of both S1 and S2 in last 166 ms |
| 25 | Percentile rank of the mean in the last 500 ms of the sum of distances between all respective body parts of S1 and S2 |
| 26 | Shortest mean of all pairwise body part distances for S2 in last 500 ms |
| 27 | Shortest median of all pairwise body part distances for S2 in last 500 ms |
| 28 | Deviation from the mean of all body part movement speed in both S1 and S2 |
| 29 | Percentile rank of sum of S1 centroid movement in last 500 ms |
